# Natural variation in chalcone isomerase defines a major locus controlling radial stem growth variation among *Populus nigra* populations

**DOI:** 10.1101/2024.10.21.618920

**Authors:** Duruflé Harold, Déjardin Annabelle, Jorge Véronique, Pégard Marie, Pilate Gilles, Rogier Odile, Sanchez Leopoldo, Segura Vincent

**Affiliations:** INRAE, ONF, BioForA, UMR 0588, 45075 Orléans, France; INRAE, URP3F, UR 0004, 86600 Lusignan, France; UMR AGAP Institut, Univ Montpellier, CIRAD, INRAE, Institut Agro, F-34398 Montpellier, France

**Keywords:** Poplar, GWAS, transcriptomics, systems biology, adaptation

## Abstract

Poplar is a promising resource for wood production and the development of lignocellulosic biomass, but currently available varieties have not been optimized for these purposes. Therefore, it is critical to investigate the genetic variability and mechanisms underlying traits that affect biomass yield. Previous studies have shown that target traits in different poplar species are complex, with a small number of genetic factors having relatively low effects compared to medium to high heritability. In this study, a systems biology approach was implemented, combining genomic, transcriptomic, and phenotypic information from a large collection of individuals from natural populations of black poplar from Western Europe. Such an approach identified a QTL and a gene, coding for chalcone isomerase (CHI), as a candidate for controlling radial growth. Additionally, analysis of the structure and diversity of traits as well as CHI gene expression revealed a high allelic fixation index, linked to the geographical origin of the natural populations under study. These findings provide insights into how adaptive traits arise, are selected, and maintained in the populations. Overall, this study contributes to enhancing the use of poplar as a valuable resource for sustainable biomass production.

## Introduction

Trees play a crucial role in mitigating climate change by sequestering carbon from the atmosphere through photosynthesis, and forest ecosystems are considered the largest terrestrial carbon sinks on Earth (Pan et al., 2011; Harris et al., 2021). The future evolution of carbon sequestration in forests relies heavily on how the growth rate and lifespan of trees respond to the changing climate (Brienen et al., 2020; Zhou, 2022). Trees keep accumulating carbon in their trunks, branches, and roots as they grow, which enables them to capture and store atmospheric carbon for several decades or possibly centuries (Green & Keenan, 2022). The major part of the tree trunk is created by the cambium, and the developing xylem constitutes a complex and dynamic system that generates wood in accordance with the seasonal cycle (Rathgeber et al., 2016). However, we still lack an integrative theory to understand growth patterns because wood formation requires the coordination of many metabolic pathways (Bryant et al., 2023).

Knowing and understanding the links between phenotypes and genetic mutations is a major challenge. Such studies have emerged for poplar, a model for tree biology, genomics, evolutionary and ecological genetics (Jansson & Douglas, 2007; Douglas, 2017). Furthermore, cultivated poplars have commercial value for peeling and veneer, lumber, paper pulp and are also used as bioenergy feedstock due to their high biomass production and favourable cell wall chemistry (Porth & El-Kassaby, 2015; Taylor et al., 2016; An et al., 2021; Abreu et al., 2022). *Populus nigra* is a deciduous tree species native to Europe, Asia and North Africa that occupies riparian ecosystems with diverse climate ranges (De Rigo et al., 2016). The genetic structure of this species in its natural distribution area is not extensively known. Yet, some studies have shown high genetic diversity within populations and low but significant genetic differentiation between river basins, suggesting high levels of gene flow in Western parts of the distribution (Smulders et al., 2008; Dewoody et al., 2015; Wójkiewicz et al., 2021). Seven ancestral genetic clusters were found in the first genome-wide genotyping study of 838 native individuals from 12 Western European river basins (Faivre-Rampant et al., 2016). However, another study of seven species showed that black poplar is highly structured with low diversity within populations (Milesi et al., 2024). These results may be due to the fact that the ecology of the species is strongly influenced by a very dynamic environment, the alluvial banks where it breeds, resulting in a complex structure (Gurnell & Petts, 2006; Alimpić et al., 2022).

Black poplar also shows a wide phenotypic diversity which can be observed on latitudinal clines such as that observed for leaf functional traits in response to drought (Viger et al., 2016) or on leaf morphology and structure (Guet et al., 2015b). Among the observable phenotypes, growth traits and wood production are considered fundamental for the adaptation and productivity of planted forests (Grattapaglia et al., 2009). For example, the biosynthetic pathway of lignin, an essential component of wood, is known to affect abiotic tolerance and growth in *Populus* (Xie et al., 2018). However, few genetic studies have been carried out on traits related to growth, and even fewer at the genomic level using the natural intraspecific diversity of trees. Genetic differentiation between natural populations of *P. trichocarpa* was found for growth and phenology, which was higher than the rather weak differentiation observed at the genome level (Evans et al., 2014; Oubida et al., 2015). This suggests that local adaptation explains patterns of variation in these traits better than genetic drift alone. The adaptive traits of poplar populations show variations depending on the local climate at their geographic origin. Using genome-wide association studies (GWAS) on provenances of *P. trichocarpa*, candidate loci underlying bud phenology and biomass have already been identified (Evans et al., 2014; Zhang et al., 2019). Based on 113 natural *P. tremula* genotypes from Sweden, a study showed significant natural variation in growth and wood-related traits and allowed the identification of genetic markers associated with these traits (Escamez et al., 2023). In this context, the OGDH enzyme (2-oxoglutarate dehydrogenase) was found to be associated with variation in tree volume and constitutes an interesting potential candidate for improving stem volume. Within the same collection of Aspen trees, a major and unique locus was also discovered. It determines the timing of bud formation and facilitates adaptation to different growing seasons and colder climates (Wang et al., 2018). A systems genetics approach in a subset of the same collection linked natural variation in lignin content and composition to responses to mechanical stimuli and nutrient availability (Luomaranta et al., 2024). Furthermore, QTLs were identified for stem and biomass traits in several mapping populations involving as parental species those typically used to generate cultivated hybrids (*P. deltoides, P. nigra* and *P. trichocarpa*). Of note, these studies reported several QTL hotspots for biomass accumulation in different environments (Rae et al., 2008, 2009; Dillen et al., 2009; Monclus et al., 2012).

Although QTL mapping studies in segregating progenies have reported QTL hotspots that explain a large part of genetic variation for growth, the QTL resolution was too limited to identify the underlying candidate genes (Rae et al., 2009). On the other hand, GWAS can make use of the rapid decay of linkage disequilibrium in forest trees (Neale & Kremer, 2011), but most studies carried out so far for growth traits have reported a limited number of loci that individually do not explain a large proportion of the genetic variance of this heritable trait (Mckown et al., 2014; Allwright et al., 2016). Many studies suggest that complex traits are controlled by multiple loci, each with rather small effects (Bradshaw & Stettler, 1995; Grattapaglia et al., 1996; Rae et al., 2007; Wade et al., 2022). To go further in understanding phenotypes and adaptation, the genomics toolbox and statistical methods as systems biology approach made available for research are constantly evolving (Pazhamala et al., 2021). The revolution comes in particular from the applications that “omics” technologies have made possible for plants such as forest trees (Plomion et al., 2016; Borthakur et al., 2022). Thus, with the progression of methodologies and the reduction in the costs of these approaches, a certain number of studies have examined at large scale of endophenotypes like transcriptomic (Chateigner et al., 2020), proteomic (Plomion et al., 2006; Castillejo et al., 2023; Teyssier et al., 2023) or even metabolomic (Rodrigues et al., 2021) in trees. Another study has advocated the use of RNAseq to jointly identify polymorphisms and quantify the transcriptomic variability across natural populations (De Wit et al., 2015). Such an approach could contribute to filling the gap between the genome and phenotypic variation for complex traits and further contribute to the explanation of their missing heritability (Maher, 2008; Chandler et al., 2014).

Here, we report an integrative approach encompassing population genetics and genomics together with transcriptomics to decipher the genetic architecture of secondary growth in natural populations of *P. nigra*. By making use of multi-omic information, we dissected a major QTL for stem radial growth identified by GWAS and pinpointed a candidate gene from the flavonoid pathway. Finally, we studied the genomic and transcriptomic diversity of the candidate gene and the phenotypic diversity across the natural populations and could show that the QTL is involved in growth differentiation, suggesting an implication in local adaptation.

## Material and methods

### Plant material and field experiments

The complete plant material and field management was previously described (Guet et al., 2015a; Gebreselassie et al., 2017). Briefly, an initial experimental design based on a total of 1,160 genotypes of *P. nigra*, representative of the species range in Western Europe, was established in two contrasting common gardens located at Orléans (France, ORL, 47°50′N 01°54′E) and Savigliano (Italy, SAV, 44°36′N 07°37′E) in 2008. In both sites, the genotypes were replicated 6 times in a randomized complete block design. A previous study, using a 12k Infinium array (Faivre-Rampant et al., 2016) was used to characterize the genetic diversity within this collection. A subset of 241 genotypes representative of the natural diversity and originating from 10 river basins was selected. Briefly, a population structure analysis on the entire collection with 5,600 SNPs and the model-based ancestry estimation in the ADMIXTURE software (Alexander et al., 2009) highlighted some introgression from the cultivated compartment (Lombardy poplar, ‘Italica’). The 241 genotypes of the present study were selected to minimize such introgression by setting a threshold of maximum 15% of the ancestral genetic group corresponding to the cultivated genotype.

### Climate data

Climatic variations across the locations of origin of the populations were analysed by applying principal component analysis on 19 annual bioclimatic variables obtained from the WorldClim dataset (Fick & Hijmans, 2017). The values used are the 30-year average (1970 to 2000) with a resolution of 1 km^2^ per grid cell obtained from the GPS location of the original natural populations. The first two principal components (PC1, 42% of explained variance, and PC2, 22% of explained variance) corresponded to the weighted precipitation and temperature variables, respectively.

### Phenotyping

We have described in detail the phenotyping of 21 traits in previous works (Chateigner et al., 2020; Wade et al., 2022). Only the circumference and the basic density of the wood were used in this study. Briefly, trees were pruned at the base after one (SAV) or two years of growth (ORL), to remove a potential cutting effect. Circumference refers to the perimeter of the stem measured at 1-m above the ground with a measuring tape. Measurements were carried out on 2-year-old trees in winter 2010-2011 at SAV and in winter 2011-2012 at ORL. Basic density was determined as previously described in (Chateigner et al., 2020). Briefly, it was measured on a piece of wood from the stem section harvested for RNA sequencing (see hereafter) following the Technical Association of Pulp and Paper Industry (TAPPI) standard test method T 258 “Basic density and moisture content of pulpwood”. For each site, the phenotypic data were analyzed with a linear mixed model to compute genotypic means adjusted for micro-environmental effects as described in (Gebreselassie et al., 2017). Before the adjustment of the model, a square root transformation was made to ensure the normality and homoscedasticity of the residuals. This transformation was only needed for circumference (not for wood basic density).

### Transcriptomic data

RNA sequencing was carried out on young differentiating xylem and cambium tissues collected in 2015 from two replicates of the 241 genotypes located in two blocks of the Orleans common garden, as described in (Chateigner et al., 2020). Sequencing reads were obtained to provide both transcriptomic and genomic data. Briefly, frozen milled tissue was used to isolate total RNA with RNeasy Plant kit (Qiagen, France), according to manufacturer’s recommendations and a treatment with DNase I (Qiagen, France) was made. Samples of young differentiating xylem and cambium tissues of the same tree were pooled in an equimolar extract before sending it for the sequencing at the POPS platform with Illumina Hiseq2000. Reads were mapped to the *P. trichocarpa* v3.0 primary transcripts (available in Phytozome 13, Goodstein et al., 2012.) using bowtie2 v2.4.1 (Langmead & Salzberg, 2012) and only transcripts with at least 1 count in 10% of the samples were kept, yielding 34,229 features. The raw count data were normalized by Trimmed Mean of M-values using the R package edgeR v3.26.4, calculated in counts per millions (CPM) and computed in log_2_(*n*+1). At the end, the CPM were fitted with a linear mixed model including batch and genetic effects to extract their genotypic Best Linear Unbiased Predictors (BLUPs). These genotypic BLUPs of transcripts were used for the rest of our analysis.

### Genotypic data

The full details of genotypic analysis have been described in (Rogier et al., 2023), including software used, data filtering criteria and final SNP selection. Briefly, genotyping data were obtained, using BWA-MEM v0.7.12 to map the reads into the *P. trichocarpa* v3.0 reference genome (available in Phytozome 13, Goodstein et al., 2012.) and the SNPs were called using 3 callers to generate a high-confidence SNP set. The 3 callers were GATK 3.1 (Van der Auwera et al., 2013), FreeBayes 0.9.20 (Garrison, 2012), and the mpileup command from SAMtools 1.3 (Li et al., 2009). Only the SNPs identified by at least 2 of the 3 callers and with less than 50% of missing values were selected. Missing values were imputed using the Fimpute v.2.2 program (Sargolzaei et al., 2014) and complementary genotyping data previously obtained with a 12 k Illumina Infinium Bead-Chip array (Faivre-Rampant et al., 2016). At the end, we obtained 878,957 SNPs and from these, 440,292 SNPs were retained for this study after filtering for a minimum allele frequency of 0.05.

### Genetic analyses

Unless otherwise stated, all analyses have been carried out with R v4.4.1 (R Core Team, 2021) under the RStudio environment (RStudio Team, 2020).

#### Partition of variance

The following bivariate mixed model was fitted to partition the variance in circumference across the two sites into between- and within-population genetic variation and their interaction with site using the R package breedR v 0.12-5 (Muñoz & Sanchez, 2024):

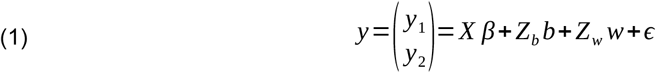

Where *y* is a vector of genotypic adjusted means for circumference in ORL and SAV, *X, Z*_*b*_ and *Z*_*w*_ are design matrices relating observations to fixed and random effects, *β* is the fixed effect of site and *b* and *w* are between and within random genetic effects. *b* and *w* follow a multivariate normal distribution with mean 0 and variances: 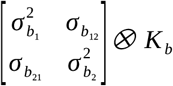 and 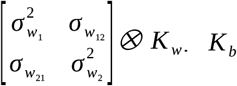 and *K*_*b*_ and *K* _*w*_ are genomic relationship matrices between and within populations. They were estimated from the full genomic relationship matrix computed with ldak software v5 (Speed et al., 2012), by averaging the kinships per population for *K*_*b*_ and setting the kinships at zero across populations for *K*_*w*_. The estimated variance-covariance parameters were then used to compute the following variance components: between and within population genetics, and between and within population genetics times environment (Itoh & Yamada, 1990).

#### Population genetics

*F*_*ST*_ was estimated using Weir and Cockerham method (Weir & Cockerham, 1984) and implemented in plink (v1.90b6.3). *Q*_*ST*_ was estimated using variance parameters from the previously described mixed-model as: 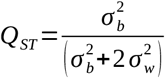

#### GWAS

GWAS was performed for circumference in each site with genotypic adjusted means and SNPs, using a linear mixed model as originally proposed by (Yu et al., 2005) and implemented in the R package MM4LMM (Laporte et al., 2022). This model included a random polygenic effect with a covariance structure defined by a genomic relationship matrix computed with the software ldak to account for linkage disequilibrium between SNPs (Speed et al., 2012). We also performed multi-locus GWAS using the multi-locus mixed-model (MLMM) approach implemented in the R package MLMM v0.1.1 (Segura et al., 2012), as well as multi-environment GWAS carried out with the MTMM approach implemented in R (Korte et al., 2012). SNPs were declared as significant according to a Bonferroni corrected threshold of 5%. Linkage disequilibrium between significantly associated SNPs was estimated in R as the squared allelic coefficient.

GWAS were also carried out using transcriptomic data (eQTL analysis) but focusing only on 2 genes of particular interest in this work, because they included significant SNPs in the GWAS. The analyses were done using both single- and multi-locus approaches, as presented for circumference.

We also looked at associations between our candidate SNP, latitude of origin and climatic data at the population level using a Pearson correlation test.

To validate the findings of present study, further tests were carried with data previously published by (Pégard et al., 2020) on a multi-parental population of *P. nigra* (factorial mating design). This dataset consisted of 629 individuals with genotypic and circumference data. We retrieved 46 SNPs within the interval [chr10:20105000, chr10:20125000] corresponding to the region of interest in the present study, and carried out association tests between these SNPs and the phenotype using a simple linear model: *y*= *X β* +*ϵ*.

## Results

This study investigates variations in stem radial growth among natural populations of black poplar using an integrative approach with multi-omics data. Phenotypic evaluations were conducted in two common garden experiments in France and Italy (Guet et al., 2015a ; Gebreselassie et al., 2017), while genotypic characterization was achieved using SNP data from RNAseq (Rogier et al., 2023). This association between phenotypic and genotypic data identified a major locus, which included two gene models annotated as a protein of unknown function (PUF) and a chalcone isomerase (CHI), respectively. To get more insights into this association, transcriptomic data from the two gene models were integrated, together with secondary traits, such as wood basic density. Finally, data from another population were also analyzed to validate the findings.

### A QTL controlling radial growth is highlighted by a genome-wide association study

We performed a GWAS for circumference using 428,836 SNPs and detected a significant signal for this trait phenotyped in Savigliano (**Fig. 1a, Fig. S1**), with a total of 18 significant SNPs, including 11 on chromosome 10 in strong linkage disequilibrium. Closer examination of this region showed that the signal is distributed over two gene models: Potri.010G212900, annotated as PUF and Potri.010G213000, annotated as CHI (**Fig. 1b**). In the MLMM approach, the whole signal vanishes out after conditioning on the top SNP, suggesting that a single allele is associated with the trait in the region (**Fig. S2**). This top SNP explained 16% of the phenotypic variation. While non-significant at the genome-wide level when considering circumference at Orleans (p-value = 0.0025, **Fig. S3**), this top SNP still explained 4% of the phenotypic variation in this common garden and its effect was in the same direction as found in Savigliano (**Fig. 1c**). Consequently, a multi-trait GWAS combining phenotypes from the two common gardens confirmed this signal but detected only a total of 7 significant SNPs (**Fig. S4**), mainly for the global effect (i.e., common to the two sites). These 7 SNPs identified in the multi-trait GWAS, are included in the 11 detected in single-trait GWAS at Savigliano and constitute our core set of candidate SNPs (**Tab. S1**). Among them, 6 are exonic (4 non-synonymous and 2 synonymous) and 1 is in 5’UTR, unsurprisingly as they come from RNAseq reads. In addition, they are all located on the CHI gene except one. It is worth mentioning that the top SNP is located in an exon of CHI gene and is predicted to be non-synonymous.

**Figure 1.**
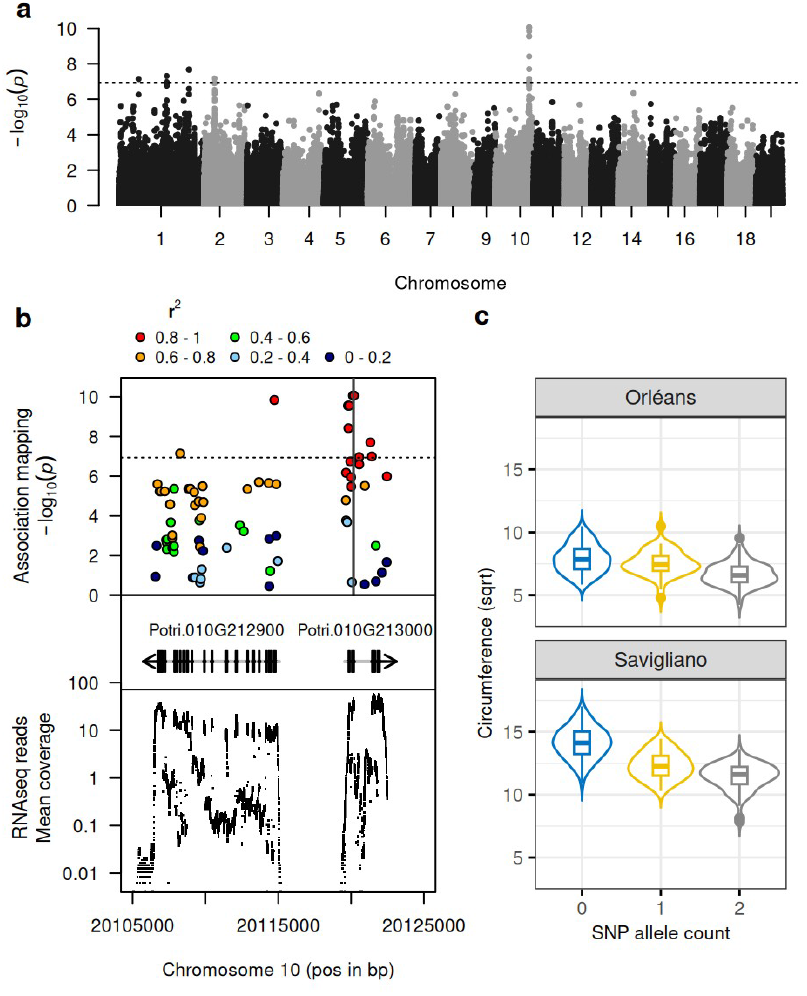
GWAS of the circumference phenotype. a) Genome-wide Manhattan plot highlighting QTL on chromosome 10 performed using a single locus mixed model and 428,836 SNPs markers from natural *P. nigra* diversity phenotype at Savigliano; b) Manhattan plot focused on SNPs with lowest p-values (colored according to their LD with the top SNP, as estimated with the squared allelic correlation coefficient r2) and concerning 2 gene models, with corresponding mean coverage of RNAseq reads across individuals; c) Box plot of the circumference in both experimental sites (transformed with a square root, sqrt), depending on the count of the alternate allele of the candidate SNP with the lowest p-value (Chr10:20120195 with reference and alternate alleles A and T, respectively).

### Gene expression sustains CHI as a candidate gene

We used the RNAseq data, generated from the xylem and cambium tissues of poplars grown at Orléans as an endophenotype to test whether the expression of our candidate genes correlated with the phenotypes and could be linked to the effect of one of them. Negative correlations were found between gene expressions and phenotypes, and their magnitude was higher for CHI than for PUF (**Fig. 2a, Fig. 2b**), with r of -0.73 (p-value < 2.2e-16) and -0.6 (p-value < 2.2e-16) for circumference evaluated in Savigliano and Orleans, respectively. When using the expression of both genes to jointly explain phenotypes, the correlation between CHI gene expression and circumference was maintained (r = -0.70, p-value < 2.2e-16 at Savigliano and r = -0.50, p-value = 3.41e-16 at Orleans) while it drastically dropped for PUF (r = 0.06, p-value = 0.372 at Savigliano and r = -0.23, p-value = 4.53e-04 at Orleans). We also made use of transcriptomic data for CHI and PUF to perform an eQTL analysis, which highlighted a strong cis control for the 2 genes (**Fig. 2c**). The fact that these two genes are close from each other and in opposite directions on the genome, together with the existence of strong LD in the region (**Fig. 1b**), generates a positive correlation between their expressions (r = 0.43, p-value = 7.1e-12). But, when focusing on the region of interest, we observed different patterns of eQTL signal between the 2 genes (**Fig. 2d**). Interestingly, the pattern of eQTL for CHI gene was similar to the one observed for circumference (**Fig. 2d, Fig. 1b**). Altogether, these results supported CHI as a candidate gene for the control of circumference variability.

**Figure 2.**
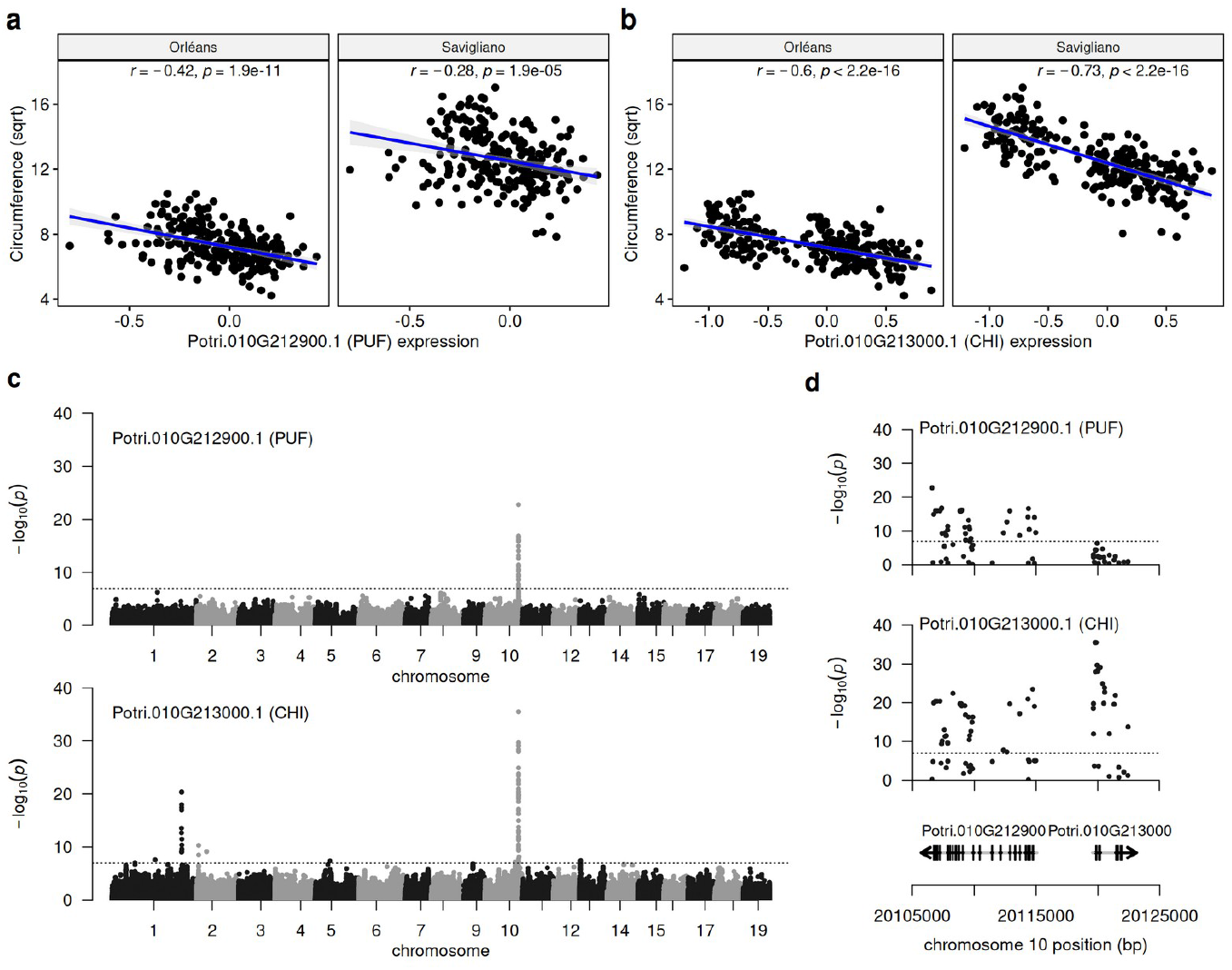
eQTL analysis sustains CHI (Potri.01G213000) as a candidate gene for the control of circumference variation. a) correlation between the circumference and the expression level of PUF primary transcript (Potri.010G212900.1) ; b) correlation between the circumference and the expression level of CHI primary transcript (Potri.010G213000). c) Manhattan plot and d) focus on the candidate region of the eQTL analysis using the variations in the expression level of the 2 primary transcripts previously highlighted as phenotypes. Circumference was transformed with a square root (sqrt). The expression level of transcripts have been standardized with a mixed linear model (see Material and methods).

### Structure of the diversity of the CHI gene highlighted by population-scale analyses

To further characterize the effect of the top SNP on the phenotypic variability, we partitioned the variance of circumference across locations into between-population and within-population genetic effects, their interaction with location, and a residual term (**Fig. 3**). This analysis showed that a large part of the phenotypic variation (35%) was due to genetic differences between populations, followed by interaction variance between genetics within populations and location (25%), genetic variance across populations (20%), and interaction variance between genetics across populations and location (17%). Interestingly, when the top SNP was included as a fixed effect in this variance partitioning model, it explained up to 24% of the total phenotypic variance, and this part of variability was mainly from the between population genetic component (**Fig. 3**, model 2). This analysis suggests that the QTL, previously identified by GWAS, is driven by differences in radial growth at the population level.

**Figure 3.**
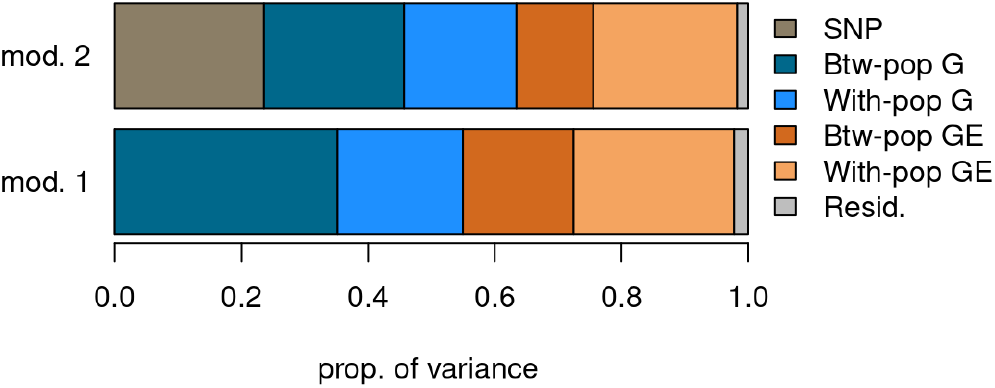
Partition of phenotypic variance for circumference across two locations using two models: Model 1 (mod.1) refers to the model of variance partition without the top SNP (Chr10:20120195), while model 2 (mod. 2) is the model that includes the top SNP as a cofactor. Btw-pop and With-pop refer to between and within population variances, while G and GE refer to genetic and genetic by environment variance, respectively.

To confirm this observation, we computed the fixation index (FST) of the 428,836 SNPs and looked at the value of the top SNP detected by the GWAS. This SNP displayed a high F_ST_ value (0.69) well above the 99^th^ percentile (0.28) of the genome-wide F_ST_ distribution (**Fig. 4a**). Such a high fixation index is due to a fixation of the reference allele in several populations mainly from the north-east of the studied area (NL, Kuhkopf, Rhin, Ticino), a fixation of the alternative allele in some population from central (Loire, Val d’Allier) and southern (Ramières) France and southern Italy (Basento), and a balanced situation in intermediate populations between these extremes (Dranse and Paglia) as well as in the population of south-western France (Adour) (**Fig. 4b**). Interestingly, such a genetic differentiation is also observed at the phenotypic level as well as at the transcriptomic level for CHI gene, as highlighted by high Q_ST_ values (**Fig. 4a**) and population differences (**Fig. 4c**). Consequently, associations between SNP and traits (**Fig. 5a**) or gene expression (**Fig. 5b**), as well as correlations between traits and gene expression (**Fig. 5c**), were high and significant when estimated at the population level, except for the trait evaluated at Orleans, which is consistent with the results obtained at the individual level (**Fig. S5**).

**Figure 4.**
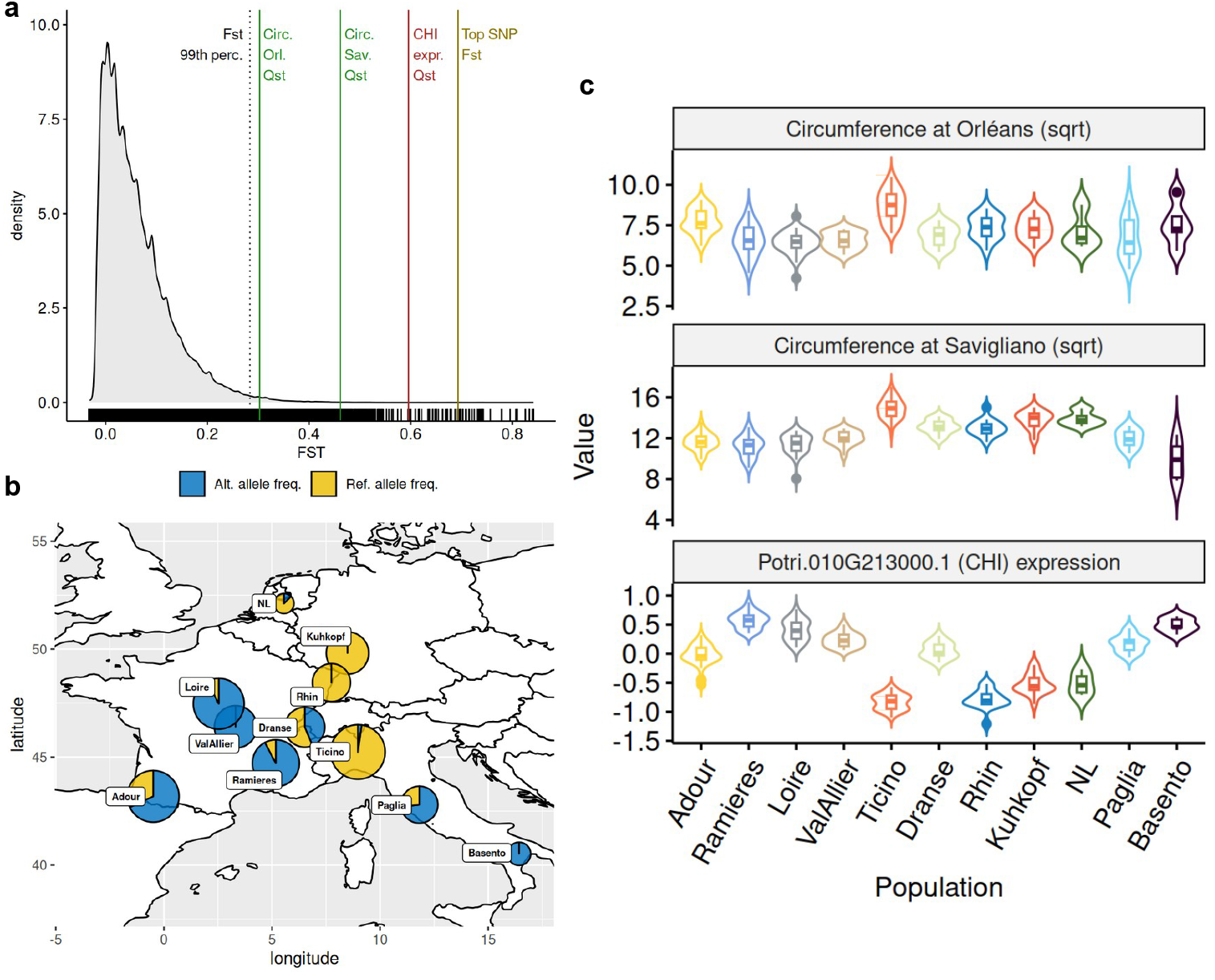
Structure of the diversity. a) Distribution of genome-wide Fst together with specific values indicated by vertical lines: 99th percentile of the distribution, top SNP (Chr10:20120195) Fst, CHI (Potri.010G213000.1) expression Qst, circumference at Orléans and Savigliano Qsts. b) Geographical origin of populations together with the distribution of alleles within each population for the top SNP (Chr10:20120195), the size of the pie is proportional to the size of the population. c) Distribution of circumferences at Orléans and Savigliano, as well as CHI (Potri.010G213000.1) expression across populations.

**Figure 5.**
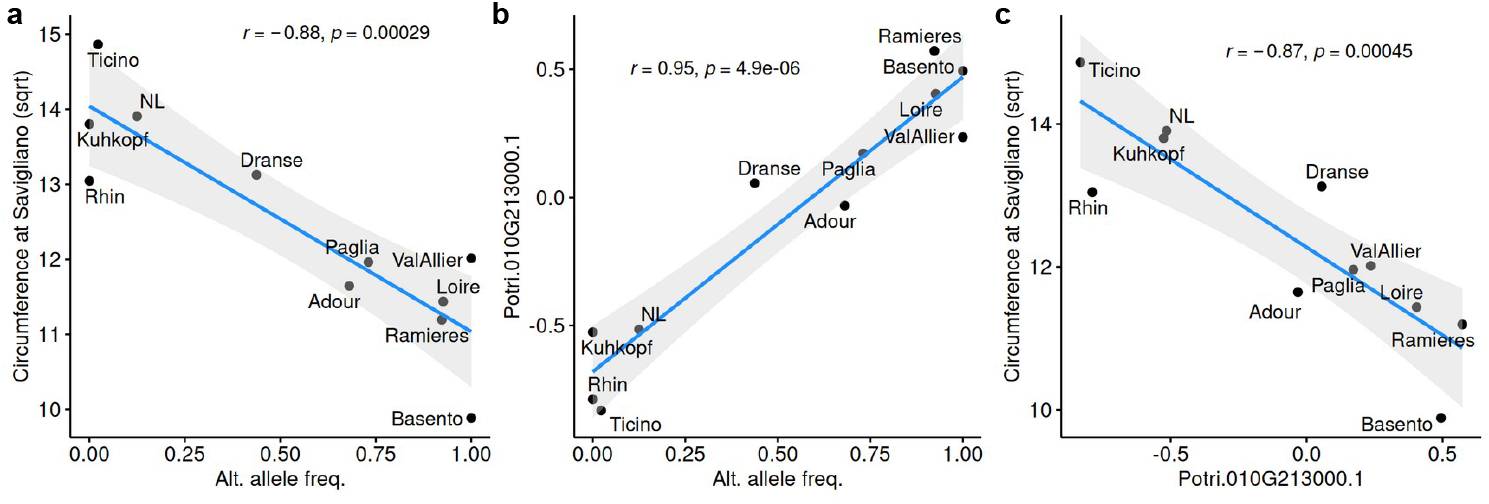
Associations at the population scale. a) Correlation between circumference at Savigliano and alternate allele frequencies for the top SNP (Chr10:20120195); b) correlation between CHI (Potri.010G213000.1) expression and alternate allele frequencies for the top SNP (Chr10:20120195); c) correlation between circumference at Savigliano and CHI (Potri.010G213000.1) expression.

### Validations and interpretations

To support our findings, we complemented our study by several analyses. First, we looked at co-localizations between the QTL detected in the present study and QTLs previously reported in the literature. Of particular interest, we found in the same genomic region a QTL previously reported by Rae et al. (2009) for several traits related to biomass production in an interspecific poplar progeny, and named poplar biomass locus 3 (PBL3). PBL3 included several QTLs for height and diameter found across multiple years, and it was delimited by two SSRs (ORPM149 and PMGC2786). We retrieved the coordinates of these markers on the *P. trichocarpa* reference genome by blasting their priming sequences. The resulting interval in bp was [17566502, 21189318] (**Fig. S6**). It thus fully includes the QTL reported here which spans the interval [20105000, 20125000] (**Fig. 1b**). Second, we retrieved data from intraspecific crosses of *P. nigra* carried out within the French breeding program and previously used and reported by (Pégard et al., 2020) for genomic prediction. From the SNP set in this previous study, we identified 46 SNPs that fell within the interval and tested associations between each of these SNPs and the phenotype circumference in a panel of 629 individuals resulting from those crosses. The most significant association (p = 2.43e-05) was found for a SNP located at 20 119 788 bp (407 bp from the top SNP) (**Fig. S7**), which was also found significant in the present study with an effect in the same direction (alternative allele associated with an increase in circumference). Finally, to provide some biological interpretation to our findings, we retrieved data on wood basic density measured on samples collected at Orléans. The top SNP displayed a significant association with wood basic density, with a positive effect of the alternate allele, which was thus opposite to the effect found for circumference (**Fig. S8a**). Similarly, a significant positive correlation was found between wood basic density and CHI gene expression while such correlation was negative for circumference (**Fig.5a, Fig.5c, Fig. S8b**).

## Discussion

We made use of circumference data collected in two common garden experiments together with transcriptome-wide SNP data to search for genetic associations between genotype and phenotype in *P. nigra*. Such analysis pinpointed a small genomic region located at the distal end of chromosome 10 which encompassed 2 gene models, of which one was annotated as a chalcone isomerase (CHI). Transcriptomic data within one of the two common gardens further supported an implication of CHI in the phenotypic variation. Because the black poplar collection was structured into subpopulations corresponding to the geographic origins of the accessions, we further focused on differences between subpopulations and found that CHI diversity is a main driver of growth differences at the subpopulation scale. Such findings suggest an implication of this gene in local adaptation. Finally, we seek to validate our results with data from previous works and found that our significant loci match a previously reported QTL hotspot for biomass accumulation in an interspecific poplar family (Rae et al., 2009). We further validated the effect of the QTL in an independent panel with a *P. nigra* pedigree from the French breeding program (Pégard et al., 2020).

The strongest effect in the GWAS was found for the phenotypic data collected in the common garden (SAV) where the genetic variability for growth was the largest. This site enabled a better expression of the phenotypic variability for growth. Unfortunately, transcriptomic evaluation was carried out in the other common garden (ORL). Consequently, it is hard to conclude on the interplay between SNP variation and gene expression to explain the variation in growth. Indeed, if we run a mediation analysis, as proposed by Sasaki et al. (2018), using phenotypic data from SAV, we cannot conclude that the expression of CHI mediates the genetic association (data not shown). While if we repeat such analysis with phenotypic data from ORL we find that the association is mediated by CHI expression, although the association with growth at ORL is not significant genome-wide. Yet, the fact that gene expression data were collected from a different site than the one in which the GWAS is significant and on trees of different ages, underlines the robustness of the results.

Another complication with the loci detected originates from the confounding effect of population structure. Indeed, the phenotype, gene expression as well as polymorphisms display a significant variability across populations which drives the correlations and associations between them. Such a situation is not ideal for association mapping, and the confounding effect attributed to population structure has to be specifically handled by the statistical model applied, usually a linear mixed model with a random polygenic effect having as covariance the genomic relationship between individuals (Yu et al., 2005). In addition, we used the program ldak to account for LD between polymorphisms in the estimation of the relationships (Speed et al., 2012), required to be independent, because our SNPs come from RNAseq and are thus clustered by genes with potentially some strong LD between neighbouring SNPs. The correction applied within the linear mixed model with such a matrix appeared to be milder than the one achieved with a regular genomic relationship matrix such as the one estimated following (VanRaden, 2008), resulting in a significant signal. Please also note that we did not consider including a fixed effect of the population structure in the model, which would inevitably clean the signal, since the phenotype is heavily structured. Such a complicated situation underlines the need to validate the association, which was achieved through two main approaches. First, we found that the detected locus falls within a QTL hotspot for biomass previously reported in several mapping populations (Rae et al., 2008, 2009; Dillen et al., 2009; Monclus et al., 2012). Second, we have shown that one of the significant SNP affects also the growth in a large collection of *P. nigra* from a breeding pedigree previously used for testing genomic prediction in black poplar (Pégard et al., 2020). While statistically significant, the effect of the SNP is lower than in the natural populations. This could potentially be explained by GxE interaction since we already found that the SNP effect is different between the two common garden experiments and the *P. nigra* pedigree from Pégard et al. (2020) was evaluated in a different location within a quite different climatic area (oceanic climate).

Another way of validating the locus would be to gain insights into the biological mechanism relating the polymorphisms to the trait through the expression of CHI. Considering the polymorphisms, we could identify four non-synonymous SNPs significantly associated with the phenotype, one in the first exon and three in the second exon of the gene, including the top SNP (**Tab. S1**). Interestingly, some of these SNPs (**Tab. S1**) are part of nucleotide triplets that map to stop codons (Chr10-20120172) or involve nucleotides very close to them (Chr10-20120195). In addition, there are two alternative transcripts for CHI gene: Potri.010G213000.3 and Potri.010G213000.2. The latter is shorter (the last exon could be missing), most likely due to the presence of SNPs linked to stop codons, implying that a variant is associated with a truncated and probably nonfunctional protein. We could thus hypothesize that one or several of these SNPs affect the enzymatic activity of CHI, for which a cellular response could be an overexpression of the gene as compensation. This would be consistent with the observed positive relationship between the most significant polymorphism and the expression of CHI (**Fig. 5b**). Also, the decrease in the enzymatic activity of CHI for individuals carrying the alternate allele of the top SNP could be consistent with the decrease in growth observed in these individuals (**Fig. 1c**). Another hypothesis to explain the negative correlation between growth and CHI expression could be a trade-off between growth and wood quality (Novaes et al., 2010). Wood basic density data provided some evidence for the effect of CHI on the trade-off between wood growth and density. CHI is a key enzyme in the flavonoid biosynthetic pathway, where it catalyzes the cyclization of a central intermediate for the production of major flavonoids such as flavanones, flavonols, and anthocyanins. Flavonoids play essential roles in defence, pigmentation, and environmental adaptation, and CHI could thus be involved in these processes. However, this metabolic pathway competes with lignin biosynthesis, as they share the common precursor p-coumaroyl-CoA (Mahon et al., 2022). Interestingly, a previous study revealed that silencing hydroxycinnamoyl-CoA shikimate/quinate hydroxycinnamoyl transferase (HCT) in Arabidopsis thaliana involves an accumulation of flavonoids and a reduction of plant growth (Besseau et al., 2007). However, this relationship between lignin and growth was later found to be unrelated to flavonoids (Li et al., 2010). Another interesting hypothesis to relate fllavonoids and growth could be the inhibitory effect of flavonoids on auxin transport, as reported in Arabidopsis thaliana (Brown et al., 2001). To test this hypothesis it would be interesting to collect phenotypic data on the roots of trees of the populations under study.

When looking at the loci diversity at the population level we found a strong differentiation, far above the genome-wide level (**Fig. 4**). Such a differentiation is thus more likely to result from differential selection than genetic drift. Of particular interest, the differentiation across natural populations was also found for CHI expression and circumference (**Fig. 4**), and we could show that the top SNP contributed mainly to the between-population component of genetic variation for growth (**Fig. 3**). As a result, highly significant correlations were found between allele frequencies, gene expression, and phenotype at the population level (**Fig. 5**). When looking at the repartition of alleles on a map representing the geographic origin of the populations, a clear North-East versus South-West differentiation appears. Such a tendency was confirmed by the significant correlation found between latitude of origin and allele frequencies (**Fig. S8a**). One could thus hypothesize that the differentiation could be related to climatic differences across Western Europe, which was confirmed by the significant correlation detected between allelic frequencies and a temperature proxy of the climate of origin (**Fig. S8b**). If we go back to the phenotypic data across populations, it’s worth noting that the southern populations with the alternate allele fixed display a lower growth and higher wood density. These data support the idea that southern populations are growing slowly as an adaptation to high summer temperatures, which ultimately underlines the adaptive relevance of the locus reported here.

This work strengthens the interest in combining transcriptomics with genomics data across large natural populations to unravel locus and genes involved in key adaptive processes such as the trade-off between growth and wood formation. Such results provide some guidance to breed future varieties of trees with improved efficiency to store carbon.

## Acknowledgements

We thank GBFOR (INRAE, Forest Genetics and Biomass Facility), https://doi.org/10.15454/1.5572308287502317E12 for management of the common garden experiment.

## Data, scripts, code, and supplementary information availability

Data are available online: https://doi.org/10.57745/DSBTGG.

Information about the RNA-seq project, from which the gene expression data come, is available in the Gene Expression Omnibus (GEO) from NCBI (accession number: GSE128482).

Raw sequences (FASTQ) are available in the Sequence Read Archive (SRA) from NCBI (accession number: SRP188754).

Information on the studied genotypes is available in the GnpIS Information System (Steinbach et al., 2013) via the FAIDARE data portal (https://urgi.versailles.inra.fr/faidare/), using the keys “Black poplar” and “POPULUS NIGRA RNASEQ PANEL” for the fields “Crops” and “Germplasm list”, respectively.

Scripts and code are available online: https://forgemia.inra.fr/vincent.segura/sybiopop_chi.git. Supplementary information is available online: https://doi.org/10.5281/zenodo.13950469

