## Supplementary materials for "Natural variation in chalcone isomerase defines a major locus controlling radial stem growth variation among *Populus nigra* populations"

**Table S1.** Table of significant SNPs with the bivariate association model across the 2 locations (top SNP marked in red).

| SNP | Chr. | Pos. | Allele 1 | Allele 2 | Gene model | Annot. | Effect | Details |
|---|---|---|---|---|---|---|---|---|
| Chr10-20114733 | 10 | 20114733 | G | A | Potri.010G212900 | exonic | synonymous | exon1:c.C105T:p.A35A |
| SNP_IGA_10_19199707 | 10 | 20119788 | A | G | Potri.010G213000 | UTR5 | NA | c.-26A>G |
| Chr10-20119817 | 10 | 20119817 | T | A | Potri.010G213000 | exonic | non synonymous | exon1:c.T4A:p.S2T |
| Chr10-20119873 | 10 | 20119873 | G | A | Potri.010G213000 | exonic | synonymous | exon1:c.G60A:p.A20A |
| Chr10-20120127 | 10 | 20120127 | C | G | Potri.010G213000 | exonic | non synonymous | exon2:c.C187G:p.Q63E |
| Chr10-20120172 | 10 | 20120172 | A | G | Potri.010G213000 | exonic | non synonymous | exon2:c.A232G:p.T78A |
| Chr10-20120195 | 10 | 20120195 | A | T | Potri.010G213000 | exonic | non synonymous | exon2:c.A255T:p.R85S |

**Table S2.** Number of genotypes per population and range of values for circumference measured in Savigliano and Orléans. The circumference was measured in mm and the presented values are transformed with a square root.

| Population | N | Circumference at Savigliano (sqrt, mm) | | | Circumference at Orléans (sqrt, mm) | | |
|---|---|---|---|---|---|---|---|
| | | min | max | mean | min | max | mean |
| Adour | 36 | 9.94 | 13.61 | 11.65 | 6.22 | 9.07 | 7.75 |
| Basento | 5 | 7.85 | 12.34 | 9.89 | 5.91 | 9.54 | 7.57 |
| Dranse | 16 | 11.92 | 14.15 | 13.13 | 5.85 | 7.82 | 6.84 |
| Kuhkopf | 19 | 11.82 | 15.00 | 13.80 | 6.06 | 8.45 | 7.34 |
| Loire | 34 | 8.04 | 12.74 | 11.44 | 4.22 | 8.05 | 6.42 |
| NL | 4 | 13.18 | 14.92 | 13.90 | 6.20 | 8.77 | 7.10 |
| Paglia | 13 | 10.54 | 13.21 | 11.96 | 4.78 | 9.06 | 6.71 |
| Ramieres | 26 | 9.10 | 13.09 | 11.20 | 4.54 | 8.40 | 6.53 |
| Rhin | 14 | 11.63 | 15.03 | 13.05 | 5.94 | 8.53 | 7.38 |
| Ticino | 44 | 12.40 | 17.03 | 14.86 | 7.02 | 10.49 | 8.70 |
| ValAllier | 19 | 10.31 | 12.96 | 12.02 | 5.68 | 7.26 | 6.55 |

**Figure S1.** QQ-plot for the single-locus mixed-model GWAS carried out for circumference at Savigliano.

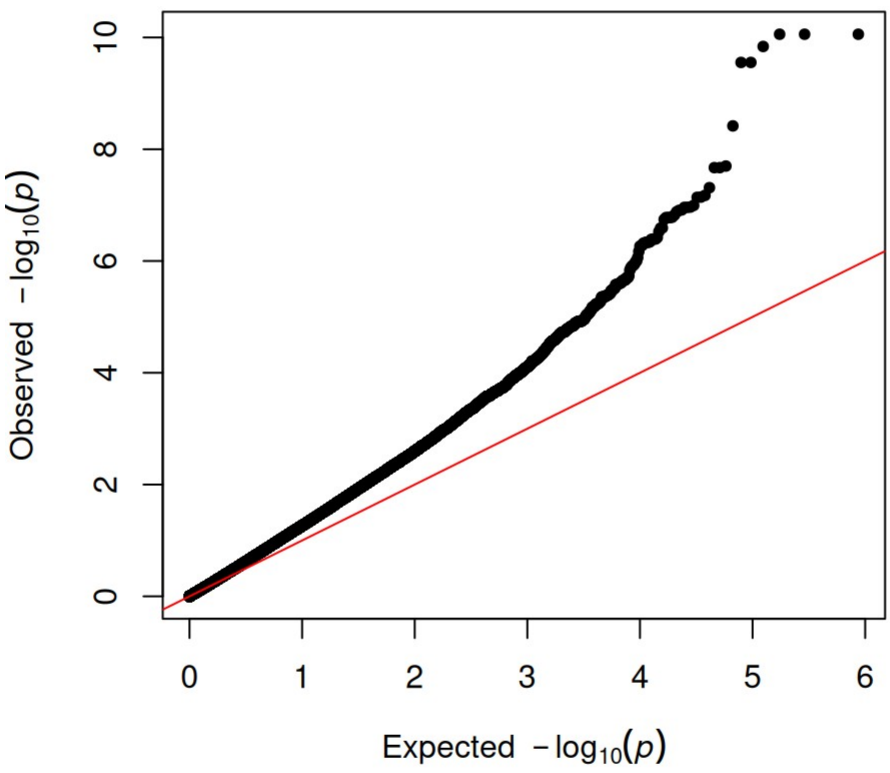

**Figure S2.** Manhattan and qq-plot for the multi-locus mixed-model GWAS carried out for circumference at Savigliano.

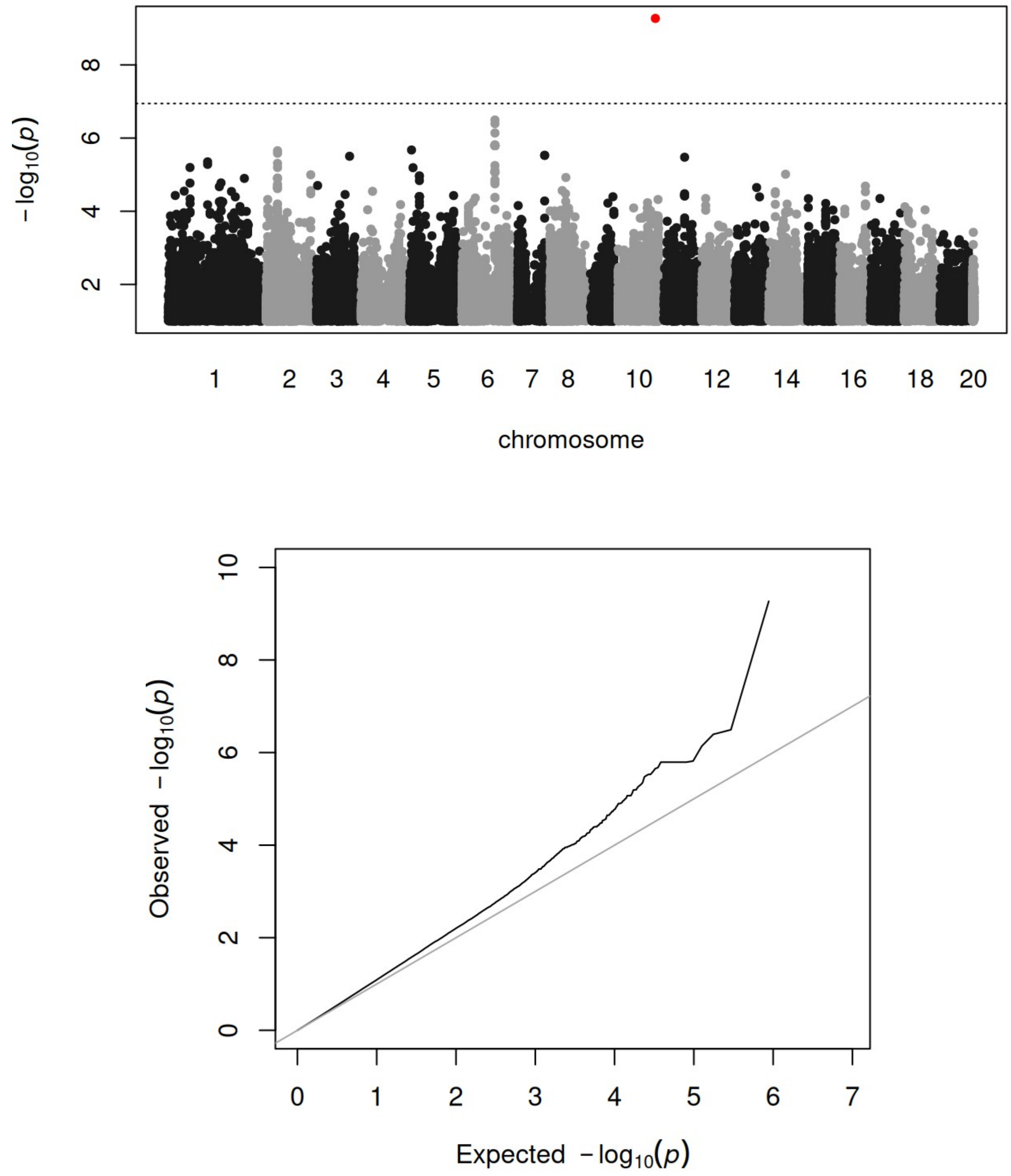

**Figure S3.** Manhattan and qq-plot for the single-locus mixed-model GWAS carried out for circumference at Orléans.

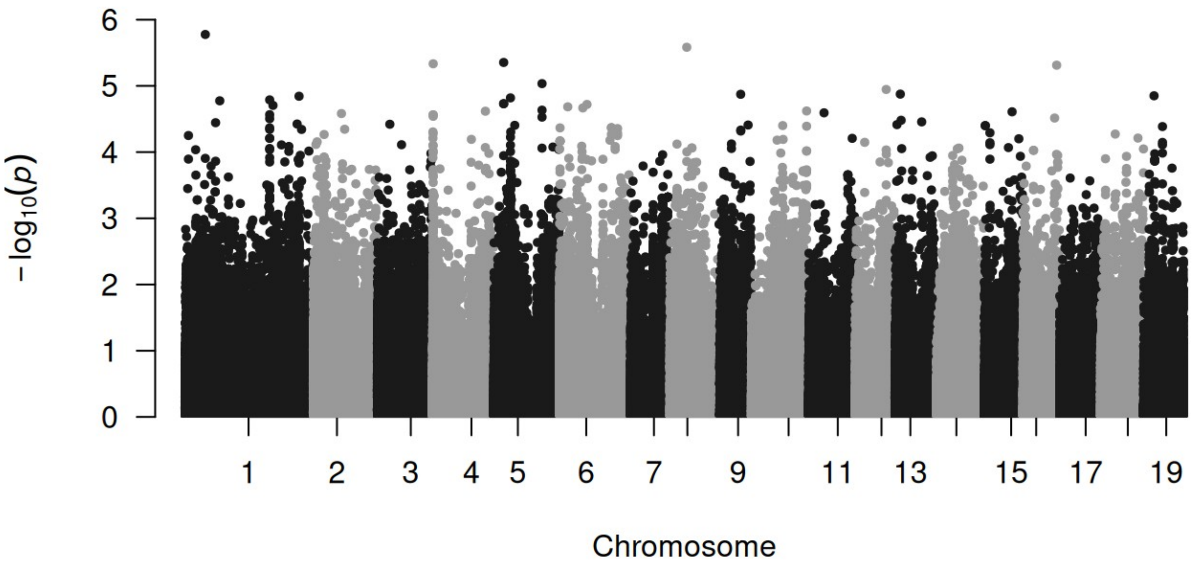


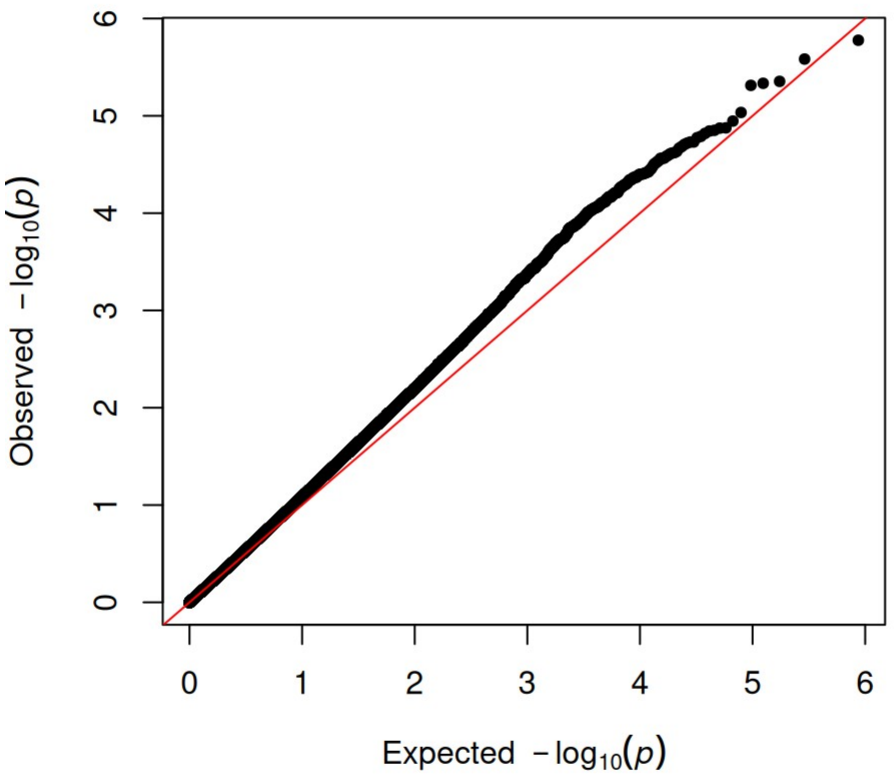

**Figure S4.** Manhattan and qq-plots for the results of the bivariate GWAS across sites for circumference. a) p-values for the global SNP effect (no GxE) ; b) p-values for the interaction effect between SNP and location.

**a.** Global SNP effect

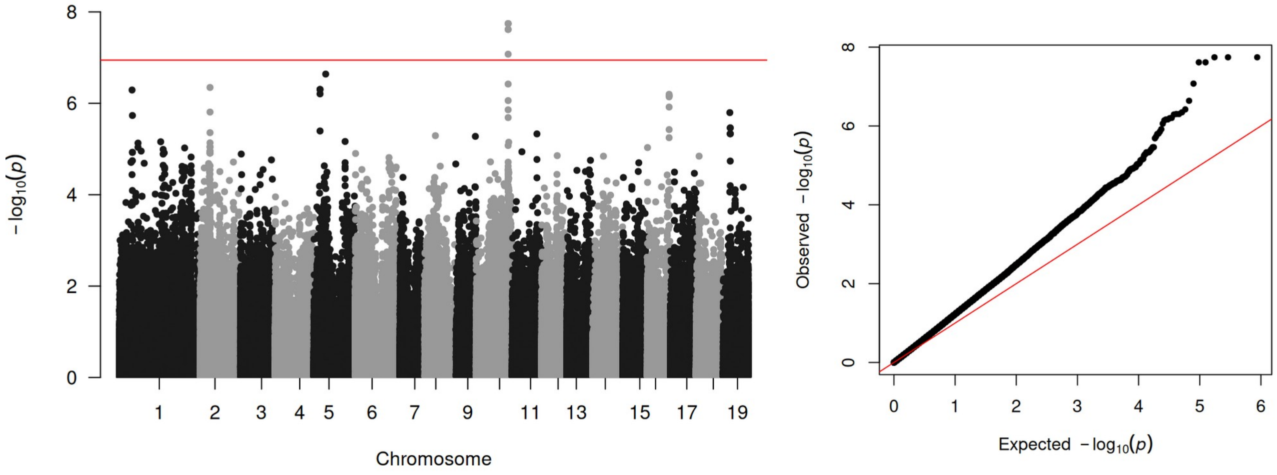


**a.** SNP by location effect

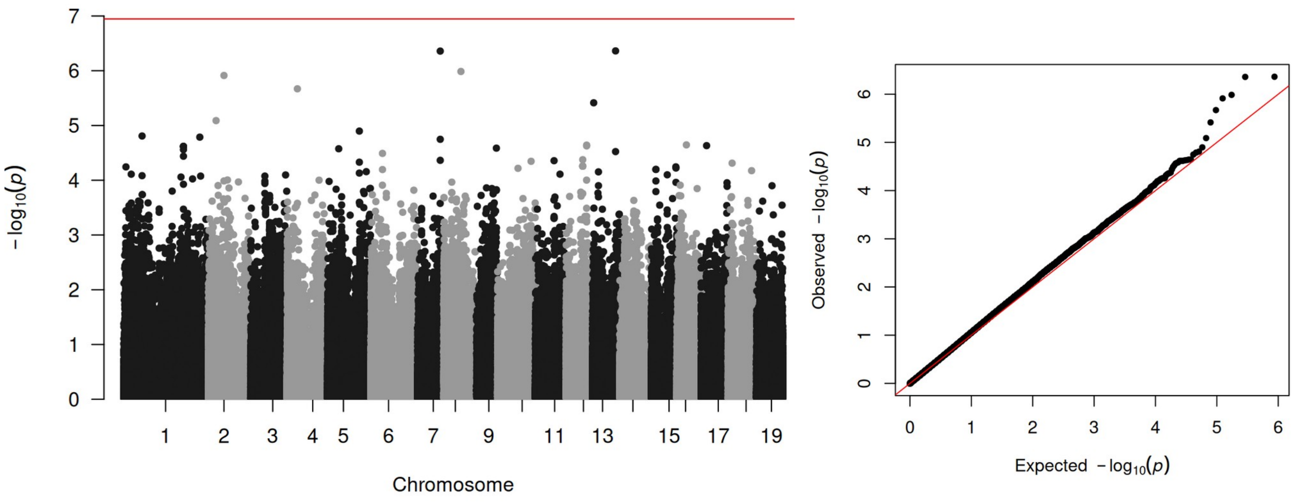

**Figure S5.** Associations at the population scale for circumference at Orléans. a) Correlation between circumference and allele frequencies for the top SNP (Chr10:20120195); b) correlation between circumference and CHI (Potri.010G213000.1) expression.

**a.**

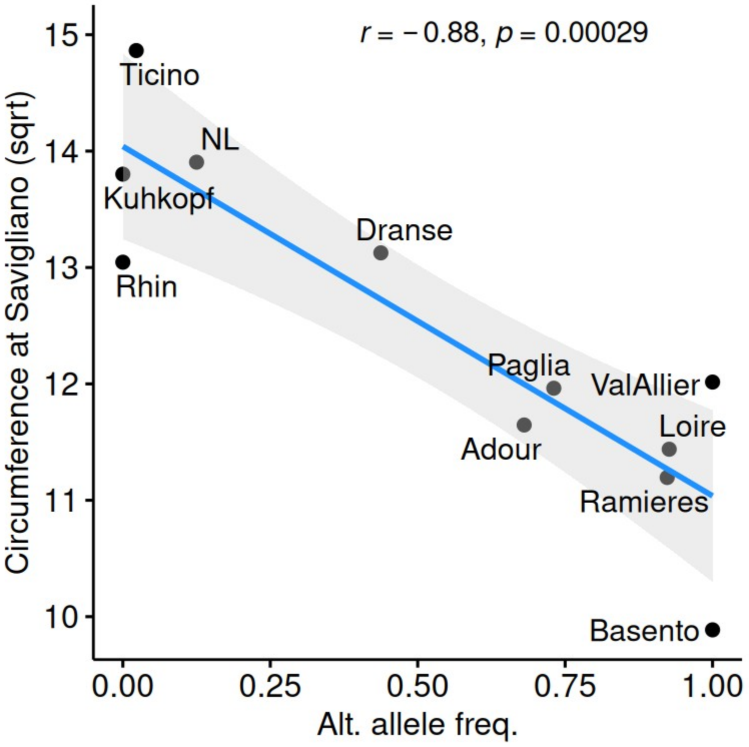


**b.**

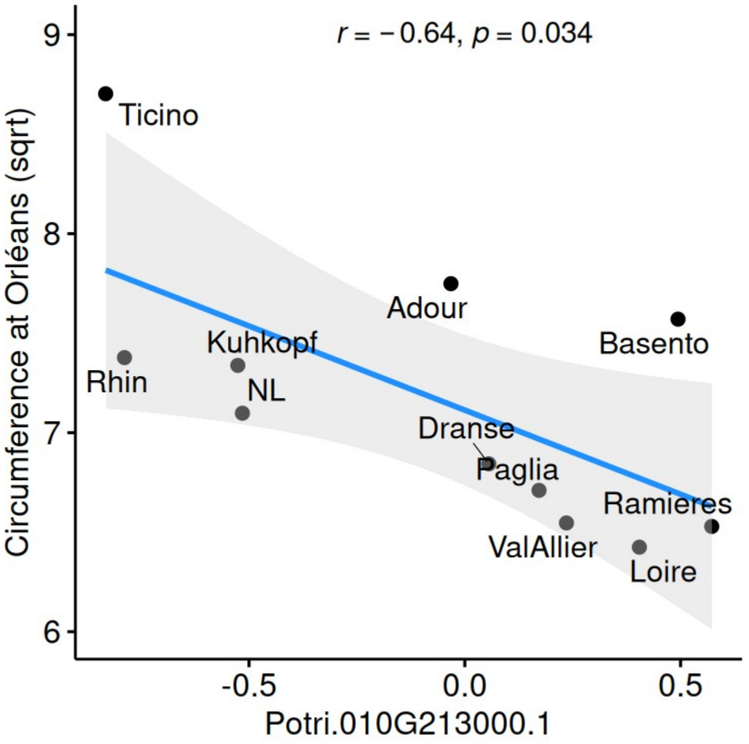

**Figure S6.** Colocalization with the poplar biomass locus reported by Rae et al. (2009).

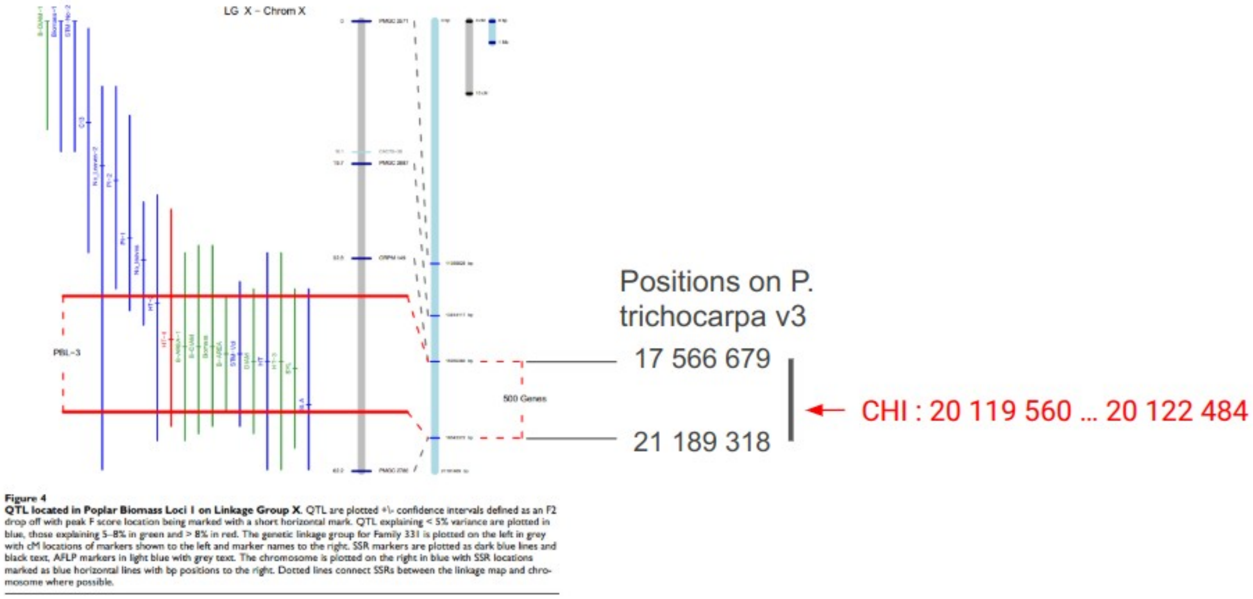

**Figure S7.** Validation of the locus effect within an external dataset (Pégard et al. 2020). Common positions between studies were retrieved in the interval [20105000 ; 20125000] and association tests were carried out with the external dataset. The most-significant SNP of such analysis ("Chr10:20119788") is presented here for its association with circumference (a) in the external dataset and (b) in the current dataset.

**a.**

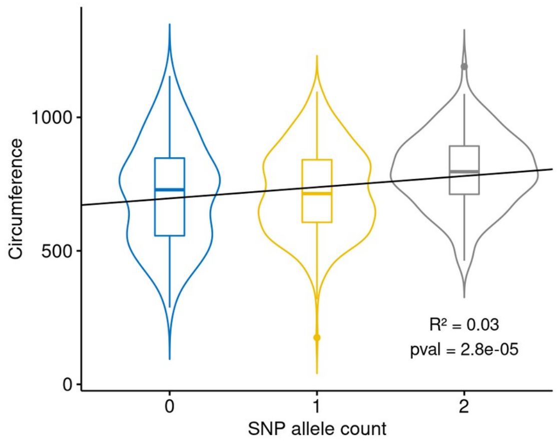


**b.**

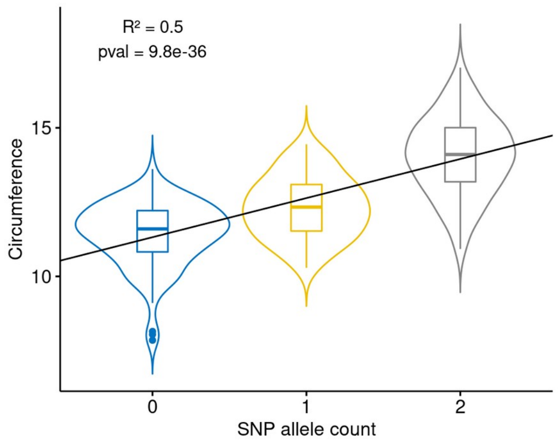

**Figure S8.** Validation of the locus effect within an external dataset (Pégard et al. 2020). Common positions between studies were retrieved in the interval [20105000 ; 20125000] and association tests were carried out with the external dataset. The most-significant SNP of such analysis ("Chr10:20119788") is presented here for its association with circumference (a) in the external dataset and (b) in the current dataset.

**a.**

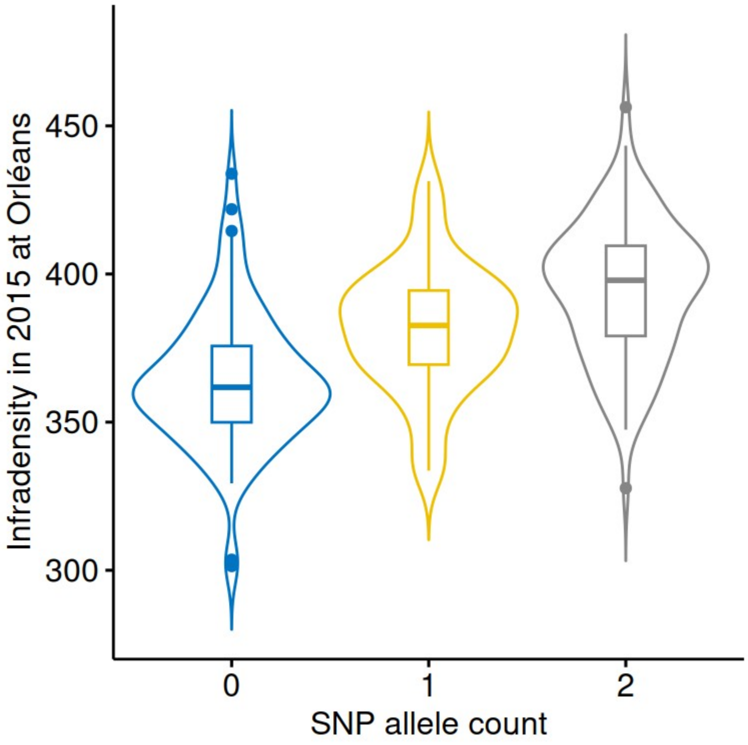


**b.**

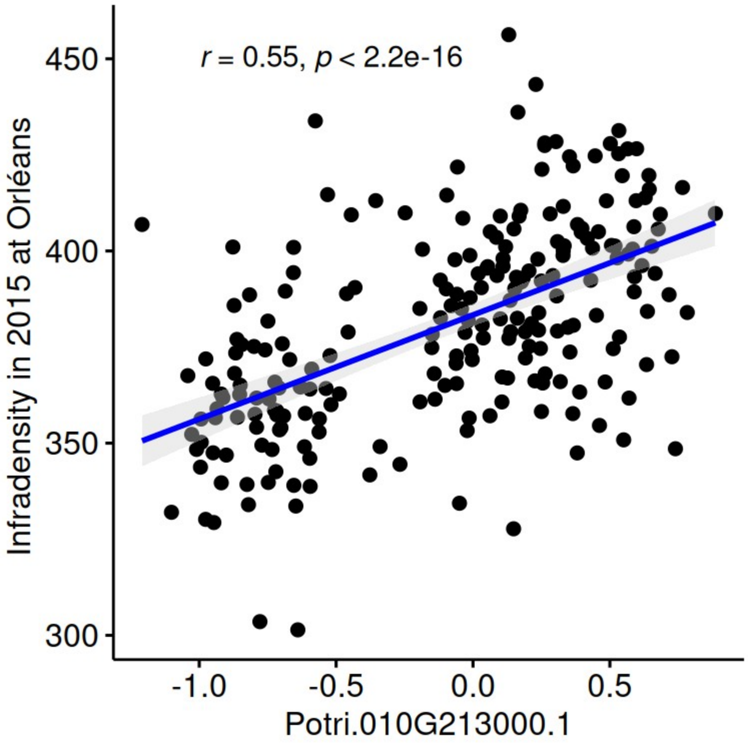
